# Impact of genetic variations on the human brain: when and where?

**DOI:** 10.1101/461814

**Authors:** Kevin Vervier, Leo Brueggeman, Jacob Michaelson

## Abstract

Genetic studies have found candidates in most of neurological and psychiatric conditions. However, it remains challenging to assess, in which brain region, and when in the brain development, a mutation is more likely to have an impact, especially for intergenic loci. Here we present, BRAVA, a publicly available resource for estimating genetic mutations spatio-temporal context on the human brain. Different contexts were observed for schizophrenia associated loci, suggesting both neurodevelopmental and late onset components.

---

Most genome-wide association study (GWAS) findings reside in non-coding and intergenic regions of the genome. Functional annotation methods have been proposed to assess how damaging a mutation can be [1,2], or what role it may have in each tissue [3,4]. The BRAINSPAN project [5] shows that gene expression in the brain strongly varies with the different regions (spatial) and development periods (temporal). To our knowledge, no existing functional annotation tool considers these extra dimensions.

We propose the Brain Variant Annotation framework (BRAVA, Fig. 1A) for estimating the spatio-temporal impact context of mutations during the brain development from fetal stage to late adulthood. Integration of brain-specific regulatory elements [6], and 3D chromosomal conformation observed in brain tissues [7] were used to capture both local and long-range contributions. We found that non-coding loci with strong statistical association in different brain regions volume (ENIGMA) also tend to have an higher RNA coverage measured in GTEx **(Supplementary Fig. 1)**.

**Figure 1.**
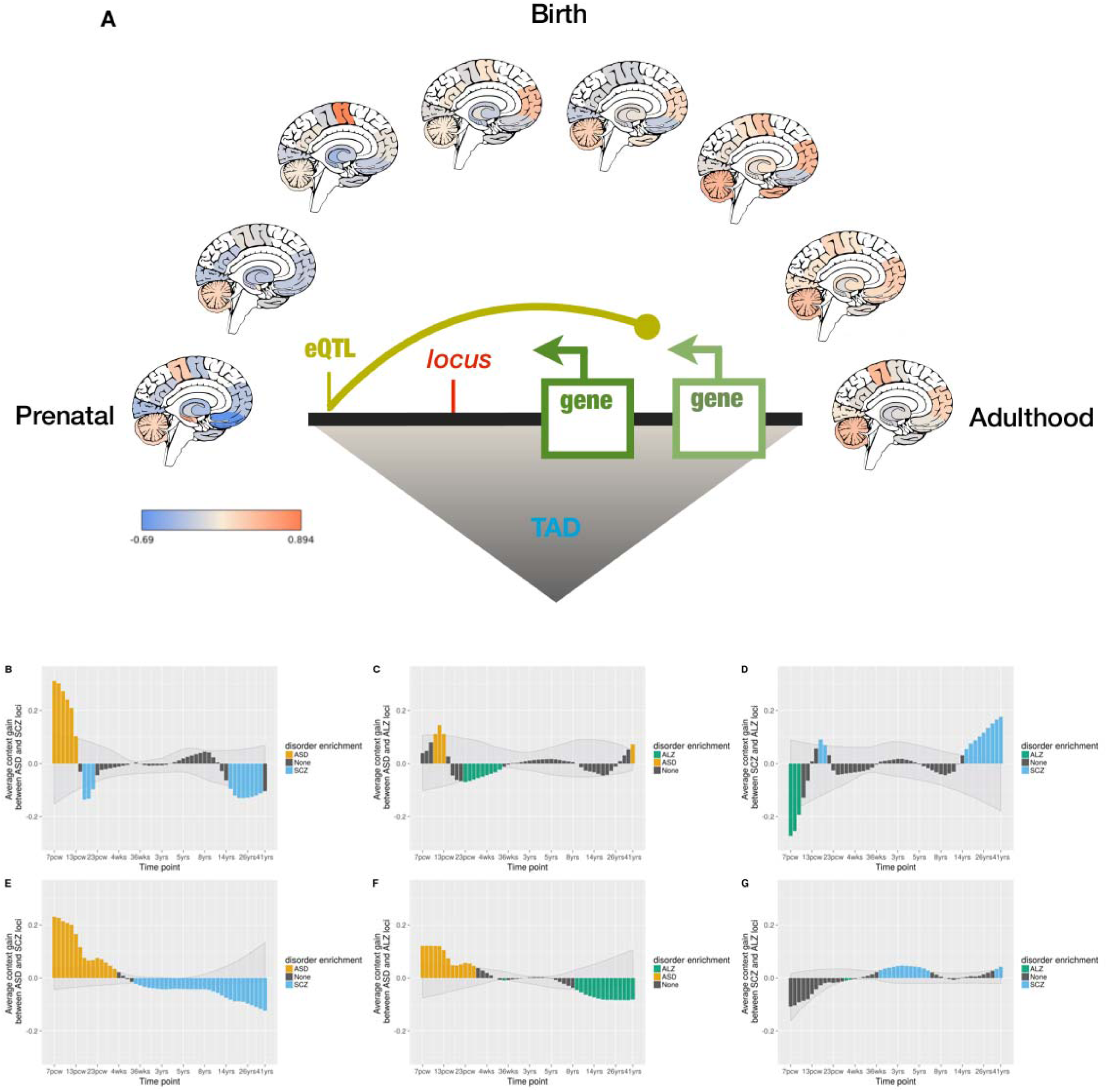
**(A)** Spatio-temporal context for genetic variations is estimated using data integration on transcriptomics, regulatory elements and chromosomal conformation information. For a given locus, the context intensity is represented in different brain regions and time points, from low value (blue) to high value (orange). Contexts for intergenic risk loci for autism (orange), schizophrenia (blue) and Alzheimer’s disease (green) are obtained by the commonly used ‘closest gene’ approach **(B,C,D)**, and by our new solution BRAVA **(E,F,G)**. Values were averaged across the 16 brain regions. Grey-shaded ribbons correspond to the null distribution estimated through 100 random permutations of genome-wide association signal. Black bars represent the absence of statistical difference between two disorders for a given time period. **(B)** Relative context enrichment for ASD and SCZ using our method. **(C)** Relative context enrichment for ASD and SCZ using the closest gene expression. **(D)** Relative context enrichment for ASD and ALZ using our method. **(E)** Relative context enrichment for ASD and ALZ using the closest gene expression. **(F)** Relative context enrichment for ALZ and SCZ using our method. (G) Relative context enrichment for ALZ and SCZ using the closest gene expression. TAD: topologically associated domain. eQTL: expression quantitative trait locus. ASD: autism spectrum disorder. SCZ: schizophrenia. ALZ: Alzheimer’s disease.

Single data sources were capturing a significant signal individually compared to a baseline defined by **(Supplementary Fig. 1-2)**. All those different data sources were then combined in a random forest framework (see Online Methods) and lead to an important improvement in terms of accuracy. Interestingly, we observed that each input dataset has a non-zero contribution to the random forest score **(Supplementary Fig. 3)**. We measured how sensitive the method is to detect specific development times for disorders with well-characterized etiology. Autism spectrum disorder is well-established as a disorder happening at both fetal [29703944] and early childhood [25007541] stages, whereas schizophrenia [2903866] and Alzheimer’s disease [9] etiologies mostly point to mechanisms occurring in adolescence or adulthood. Non-coding variations context was compared between these three disorders in **Figure 1**, and we found overall improvement in predicted patterns for our method **(Fig. 1E,F,G)**, compared to the standard approach consisting in looking for the closest gene expression profile (Fig. 1B,C,D). Especially, autism risk loci are consistently enriched in fetal brain development, over later onset disorders. We also validated the sensitivity of the method to detect known region-specific context. Studies like ENIGMA (cite) provide an unprecedented combination of genetics and neuroimaging data, allowing to consider potential interactions. Especially, we focus on two regions present in both ENIGMA and BRAINSPAN, namely the hippocampus and the amygdala. For these regions, we demonstrate that the stronger associated loci also correlate with a region-specific enrichment in BRAVA estimated context **(Supplementary Fig. 4A-B)**. Finally, we focus on the major histocompatibility complex (MHC) region strongly associated with schizophrenia risk (cite). Interestingly, the two principal risk loci **(Fig. 2A)** have been found to be statistically independent, but there is limited information about actual etiology differences. BRAVA’s context matrices suggest that C4A region **(Fig. 2C)** has a stronger effect for time points spanning from adolescence to adulthood. However, rs13194504 **(Fig. 2B)** estimated context shows a peak during fetal development across all the brain regions. BRAVA score decomposition for rs13194504 reveals a major contribution (20% of the signal) for gene GPX6 involved in the glutathione synthesis, which is a known mechanism implicated in schizophrenia [PMID: 16410648] and recently associated with fetal brain development, ethanol exposure [PMC5751199], and depression in pregnancy [27174401]. In summary, we demonstrated that our new method, BRAVA, can efficiently estimate of spatiotemporal context in the human brain, especially for non-coding loci. In particular, we foresee BRAVA application to improve the functional characterization of disease-causing variants.

**Figure 2.**
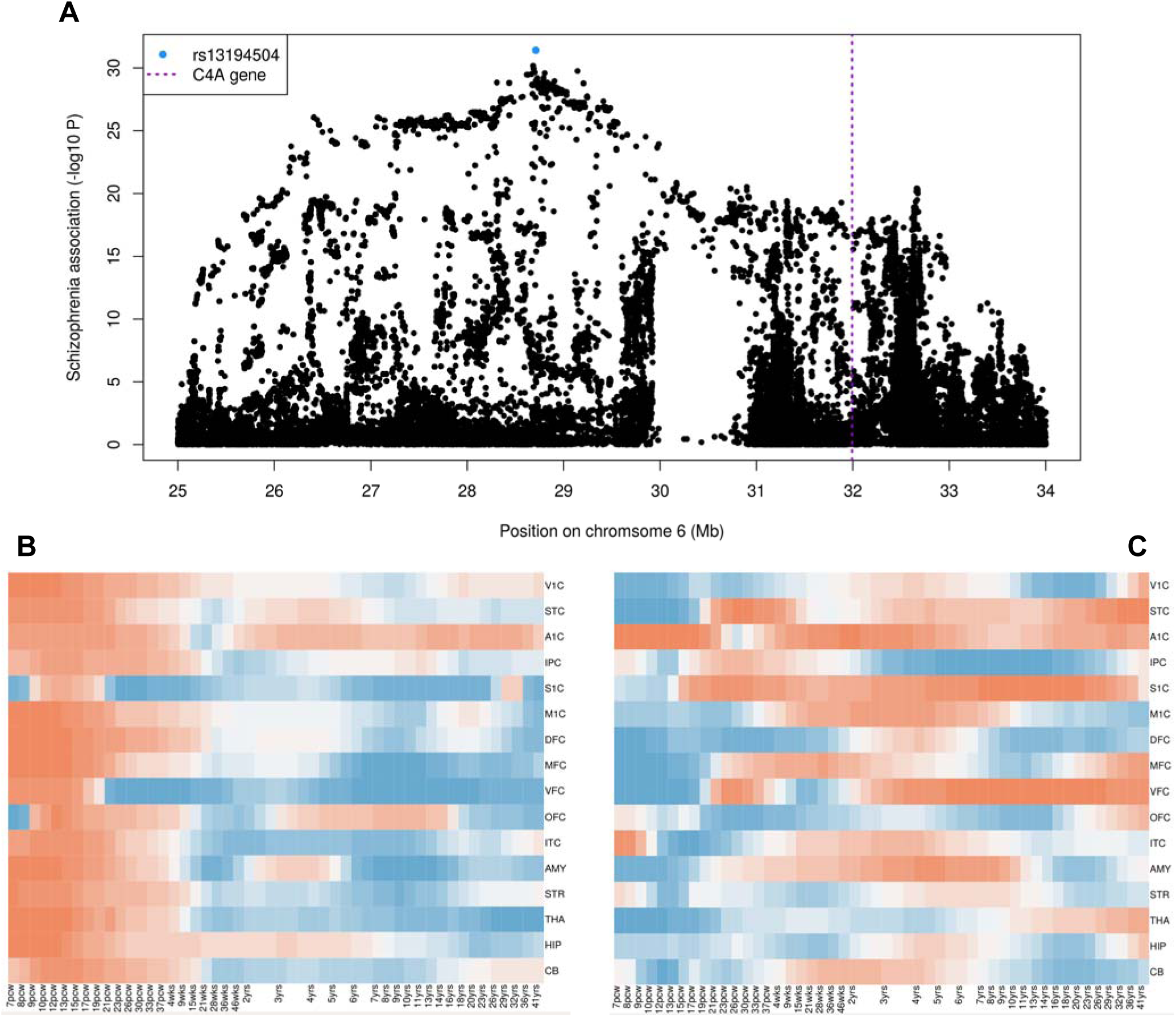
**(A)** schizophrenia-associated loci in the major histocompatibility complex (MHC) region: non-coding *rs13194504* (blue) and *C4A* gene (purple). The height of each point represents the statistical strength (−log10 *P*) of association with schizophrenia measured in the Psychiatric Genomics Consortium. **(B)** BRAVA estimated context for *rs13194504* (*chr6:31,992,884*) **(C)** BRAVA estimated context for *C4A* gene (locus at the center, *chr6:31,992,884*). Positive context values are in orange, and blue cells correspond to negative context values.

## Methods

Methods, including statements of data availability and statistical framework description, are available in the online version.

### Online Methods

#### BRAVA for spatiotemporal context estimation

We provided an R package to automatically estimate the spatiotemporal context for any genetic variation (http://github.com/kevinVervier/BRAVA).

#### Brain transcriptome data (BRAINSPAN)

Gene expression was obtained in tissues from 16 brain regions (see list in Supplementary Table 1). Individuals’ age spans between 7 post-conception weeks (fetal stage) to 41 years old. Missing data were imputed using smoothing spline (Chambers & Hastie (1992)). For each gene, expression was corrected for the average expression across the XYZ considered genes, then z-scaled across time points and brain regions (average value in a context matrix is zero, and standard deviation is one). Finally, we averaged all the gene expression matrices to estimate the background trend in expression and subtracted it to each matrix.

#### ENIGMA genome-wide association studies

Summary statistics from the ENIGMA2 project (Hibar, 2015) were downloaded on enigma.ini.usc.edu. Associations (P-values) for hippocampus and amygdala volumes were used during the model training and validation.

#### Chromosomal conformation in human brain

Data from Gerschwind, 2016 from cortical plate cells (GSE77565). Topological associated domains (TAD) and interchromosomal contact probabilities were filtered if values were lower than the 75^th^ percentile, following Gerschwind et al. guidelines.

#### Expression quantitative trait loci in brain tissues

Brain eQTLs from Gene-Tissue expression project (GTEx) were downloaded from the version v6. Statistically significant loci were considered if they were detected in at least one of the GTEx brain tissues.

#### Psychiatric Genetics Consortium data

Genome-wide summary statistics for both schizophrenia and autism spectrum disorder were downloaded on the PGC portal (med.unc.edu/pgc). The 10 independent regions with the lower P-values were used in the analysis (see lists in Supplementary Tables 2 for schizophrenia and 3 for autism).

#### Genetic association with Alzheimer’s disease

International Genomics of Alzheimer’s Project (IGAP) is a large two-stage study based upon genome-wide association studies (GWAS) on individuals of European ancestry. In stage 1, IGAP used genotyped and imputed data on 7,055,881 single nucleotide polymorphisms (SNPs) to meta-analyse four previously-published GWAS datasets consisting of 17,008 Alzheimer’s disease cases and 37,154 controls (The European Alzheimer’s disease Initiative - EADI the Alzheimer Disease Genetics Consortium - ADGC The Cohorts for Heart and Aging Research in Genomic Epidemiology consortium - CHARGE The Genetic and Environmental Risk in AD consortium - GERAD). In stage 2, 11,632 SNPs were genotyped and tested for association in an independent set of 8,572 Alzheimer’s disease cases and 11,312 controls. Finally, a meta-analysis was performed combining results from stages 1 & 2. The 10 independent regions with the lower P-values were used in the analysis (see list in Supplementary Table 4).

#### Reference genome

Analysis performed in this study used the hg19 reference genome.

#### Benchmark of various data sources on context estimation

We considered all ENIGMA GWAS SNPs and mapped them to their nearest gene. Only intergenic SNPs with distance > 10000bp were selected for further analysis (*n*=974,045). In order to compare the evaluated methods, we defined a criterion based on the ability of a prediction to recapitulate the actual association strength, in terms of p-value. Spearman rank correlation was applied to estimate the correlation, and the negative log10 of the p-value was used to compare the different methods.

#### Machine-learning algorithm

After observing each separated data source ability to estimate context, we evaluated a weighted combination of these sources. BRAVA model relies on the Random forest algorithm (Breiman) to define the relative importance of each separate data source in predicting context for adult in hippocampal region from ENIGMA. Raw Gini importance values estimated for each feature were normalized as proportions, and later used as weights in the final spatio-temporal context estimation.

## Acknowledgements

We thank the International Genomics of Alzheimer’s Project (IGAP) for providing summary results data for these analyses. The investigators within IGAP contributed to the design and implementation of IGAP and/or provided data but did not participate in analysis or writing of this report. IGAP was made possible by the generous participation of the control subjects, the patients, and their families. The i-Select chips was funded by the French National Foundation on Alzheimer’s disease and related disorders. EADI was supported by the LABEX (laboratory of excellence program investment for the future) DISTALZ grant, Inserm, Institut Pasteur de Lille, Universite de Lille 2 and the Lille University Hospital. GERAD was supported by the Medical Research Council (Grant n° 503480), Alzheimer’s Research UK (Grant n° 503176), the Wellcome Trust (Grant n° 082604/2/07/Z) and German Federal Ministry of Education and Research (BMBF): Competence Network Dementia (CND) grant n° 01GI0102, 01GI0711, 01GI0420. CHARGE was partly supported by the NIH/NIA grant R01 AG033193 and the NIA AG081220 and AGES contract N01-AG-12100, the NHLBI grant R01 HL105756, the Icelandic Heart Association, and the Erasmus Medical Center and Erasmus University. ADGC was supported by the NIH/NIA grants: U01 AG032984, U24 AG021886, U01 AG016976, and the Alzheimer’s Association grant ADGC-10-196728.

## SUPPLEMENTARY MATERIALS

**Supplementary Figure 1:**
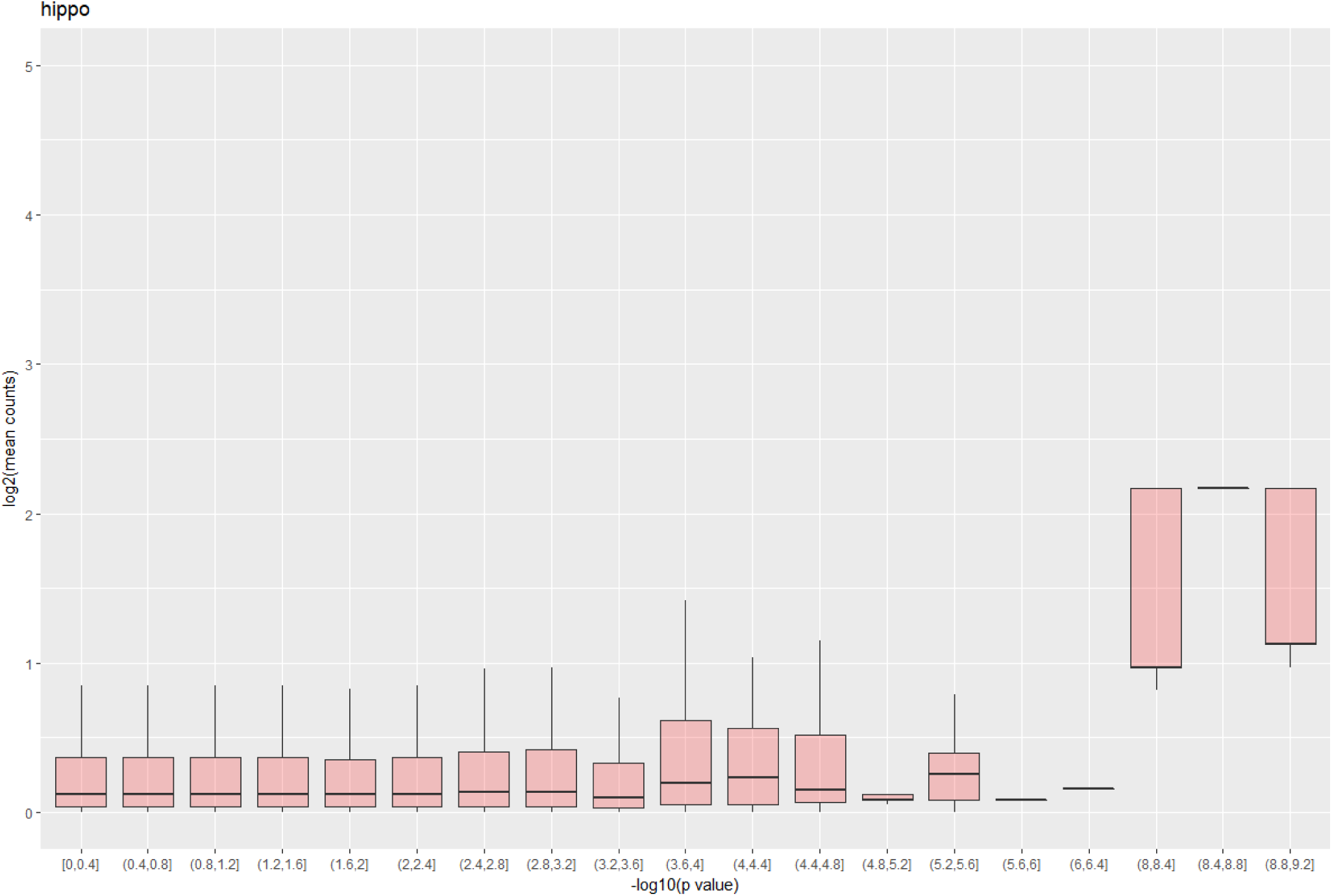
non-coding loci with strong statistical association in hippocampus volume from ENIGMA GWAS have higher RNA coverage measured in postmortem brain tissues from GTEx. x-axis: GWAS association as log10 transformed p-value. y-axis: RNA coverage as log2 transformed value.

**Supplementary Figure 2:**
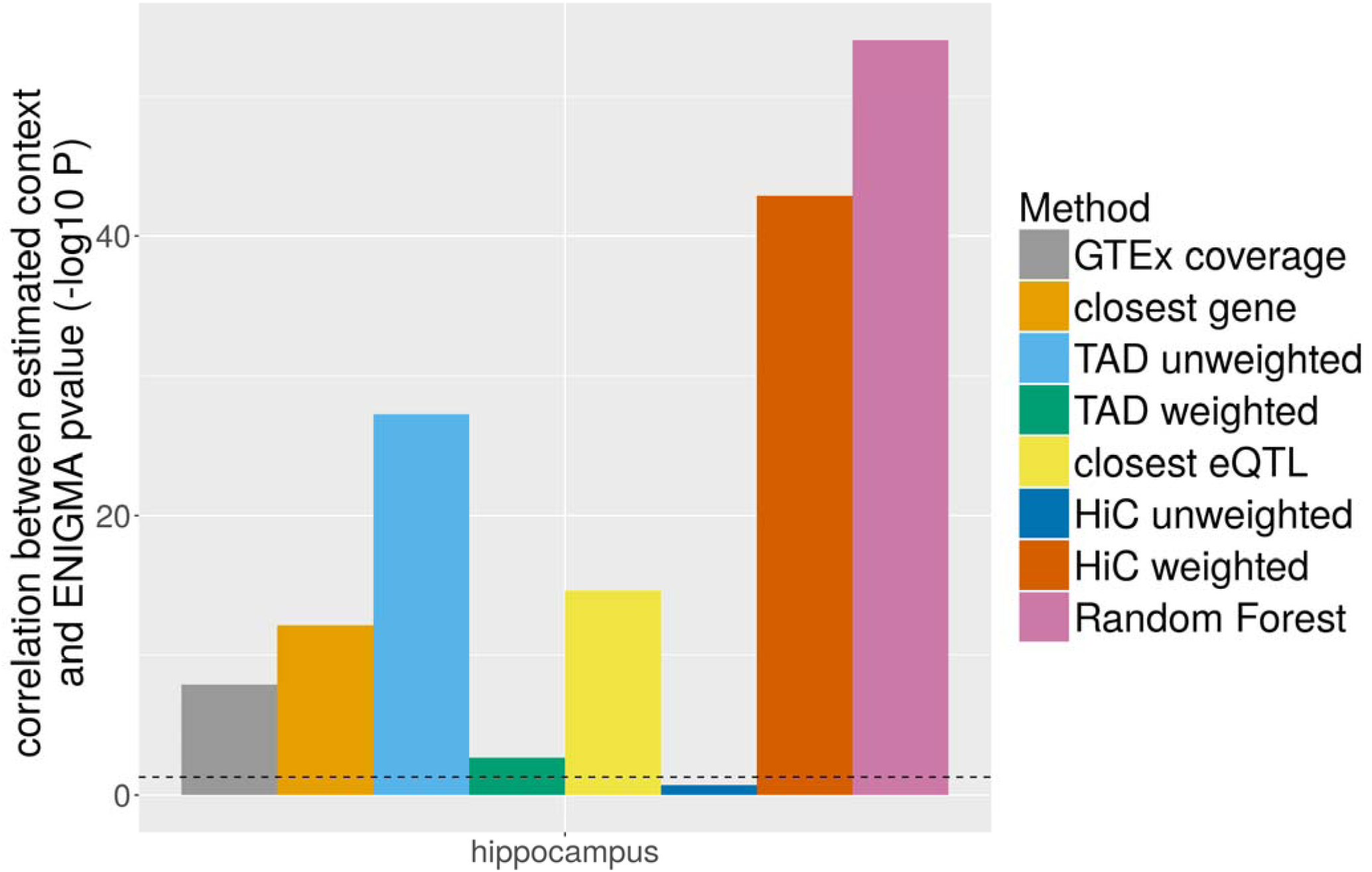
Spearman correlation between estimated variant context and ENIGMA GWAS pvalue for hippocampus volume, estimated using different data sources. Random Forest model (pink) combines all the other sources as features in a non-linear multivariate fashion.

**Supplementary Figure 3:**
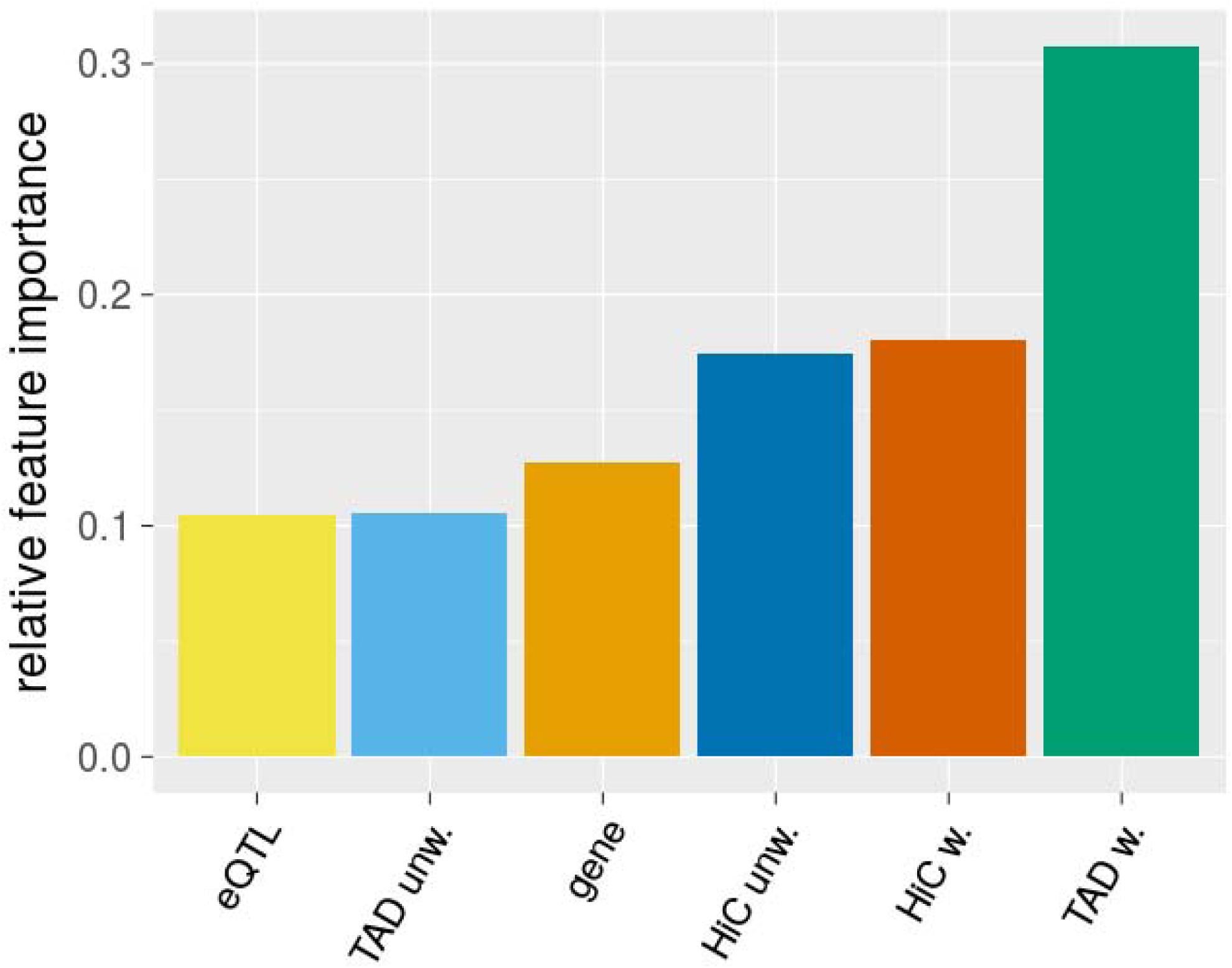
Importance of each data source in the random forest model estimated as relative Gini index measure.

**Supplementary Figure 4:**
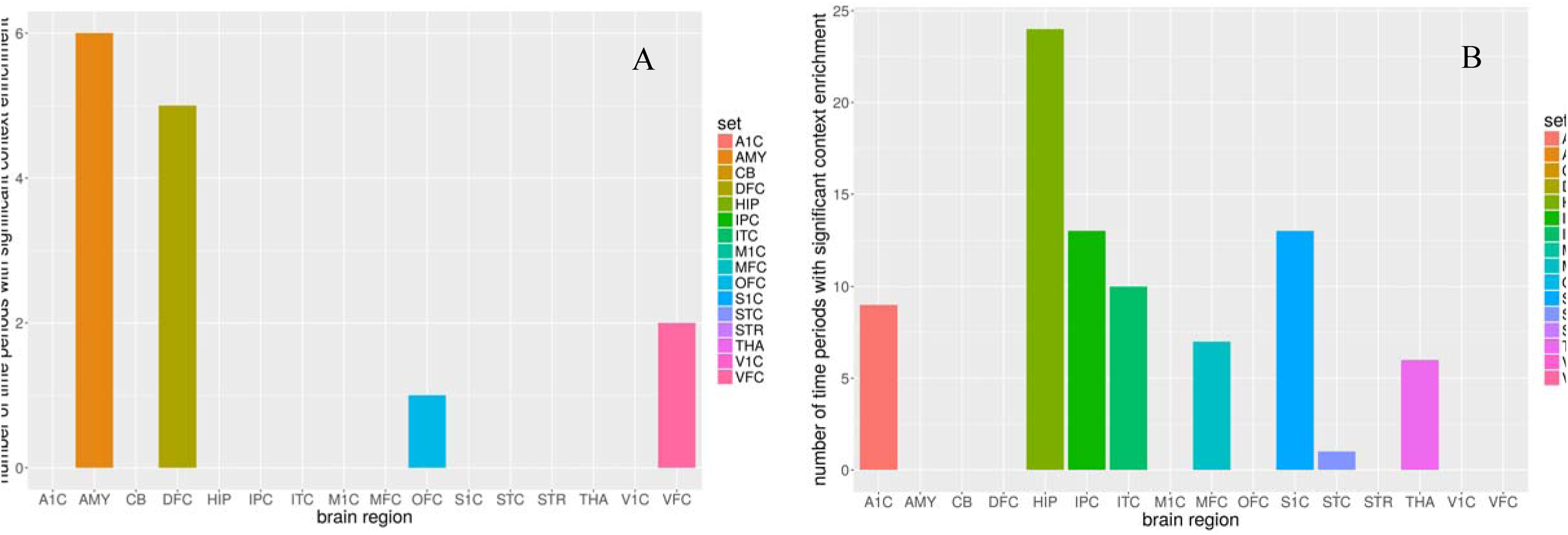
regional enrichment for amygdala (A) and hippocampus (B) as the number of developmental time points with significant enrichment over random genomic loci.

Supplementary Tables […]

